# Species-Dependent Limitations of Murine Myocardium for Investigating the Combined Inotropic Effects of SERCA2a Stimulation and Myofilament Ca^2+^ Sensitization

**DOI:** 10.64898/2026.10.30.762932

**Authors:** Francesca Misitano, Natasha Giacomelli, Claudia Maniezzi, Marica Dente, Marina Scardigli, Corrado Poggesi, Josè Manuel Pioner, Gabriella Malfatto, Antonio Zaza, Cecilia Ferrantini, Francesco Lodola

## Abstract

Impaired sarcoplasmic reticulum Ca^2+^ handling and myofilament function both contribute to contractile dysfunction in heart failure. A previous study in failing human myocardium showed that combining the SERCA2a activator istaroxime (ISTA) with the Ca^2+^ sensitizer levosimendan (LEVO) produces greater positive inotropy than either drug alone. However, human multicellular preparations did not allow comprehensive investigation of the underlying cellular mechanisms. We therefore assessed contractile performance and Ca^2+^ dynamics in murine multicellular myocardial preparations and ventricular cardiomyocytes treated with ISTA (20 µM), LEVO (0.5 µM), or their combination. ISTA improved developed force, contraction–relaxation kinetics, and mechanical restitution. These effects were associated with increased Ca^2+^ transient amplitude, accelerated Ca^2+^ rise and decay, and enhanced sarcoplasmic reticulum reloading. Adding LEVO produced no further mechanical benefit and only minor changes in Ca^2+^ handling. LEVO alone elicited heterogeneous contractile responses and increased myofilament Ca^2+^ sensitivity in only a minority of cardiomyocytes, while consistently accelerating Ca^2+^ transient decay. This effect was reproduced by the phosphodiesterase-3 inhibitor cilostamide, after which LEVO caused no significant additional change, consistent with a contribution of downstream cAMP–PKA signaling to the acceleration of Ca^2+^ removal. Thus, limited murine responsiveness to LEVO-mediated myofilament sensitization prevented mechanistic reconstruction of the additive inotropic effect observed in human myocardium, highlighting a species-dependent limitation of the mouse model for studying this combined therapy.

## 1. Introduction

Heart failure (HF) is a major and growing global health burden characterized by the inability of the heart to adequately meet the metabolic demands of the body. While chronic HF therapies primarily target neurohumoral activation and adverse cardiac remodeling, impaired myocardial contractile and relaxation function remains a major therapeutic challenge [1,2]. In patients with acute decompensated HF, characterized by reduced cardiac output and systemic hypoperfusion, positive inotropic agents may be required to restore adequate hemodynamic function [3]. However, currently available inotropes, agonists of PKA signaling in particular, act by targeting multiple effectors of excitation-contraction (EC) coupling, an approach associated with increased arrhythmic risk, higher energetic demand and adverse long-term outcomes [1,3,4].

An alternative strategy is to selectively modulate individual components of EC-coupling [1,4]. Among these approaches, direct activation of sarcomere function through Ca^2+^ sensitizers improves systolic performance without increasing cytosolic Ca^2+^ [4,5]; on the other hand, enhanced myofilament activation might impair relaxation and thereby compromise diastolic function [6]. Conversely, restoration of sarcoplasmic reticulum (SR) Ca^2+^ uptake through SERCA2a stimulation improves intracellular Ca^2+^ compartmentalization, accelerating relaxation while potentially reducing arrhythmogenic Ca^2+^ dysregulation [4,7–11]. Because both SR dysfunction and impaired myofilament performance contribute to the contractile deficit of HF [12,13], their simultaneous pharmacological targeting represents an attractive strategy to achieve balanced ino-lusitropic stimulation while limiting the drawbacks associated with conventional inotropic therapies.

In line with this rationale, preliminary evidence from our laboratory provided evidence that combined treatment with the SERCA2a activator istaroxime (ISTA) and the Ca^2+^ sensitizer levosimendan (LEVO) may improve inotropy of failing human myocardial trabeculae more than either drug alone. While these findings provide proof of principle for the combined approach, the preparation used does not allow comprehensive investigation of the mechanisms underlying this promising functional interaction.

The present study was therefore designed to dissect the cellular basis of the interaction between ISTA and LEVO using murine multicellular myocardial preparations and ventricular cardiomyocytes. By studying the effects of each agent alone and in combination on myocardial force development, relaxation, and intracellular Ca^2+^ handling, we aimed to explore the mechanisms underlying the additive functional effects previously observed in human myocardium.

## 2. Results

To investigate the cellular mechanisms underlying the functional interaction previously observed in human myocardium, we performed complementary experiments in multicellular myocardial preparations and in isolated murine ventricular cardiomyocytes. ISTA was used at 20 µM, a concentration previously characterized in the murine model as producing predominantly SERCA2a-dependent effects, with only a limited contribution from Na^+^/K^+^-ATPase inhibition [14]. LEVO was tested at 0.5 µM, a concentration at the ceiling of the clinically relevant range [15].

In multicellular myocardial preparations, a sequential experimental protocol was adopted: tissues were first exposed to ISTA, followed by the addition of LEVO while maintaining ISTA exposure. In isolated cardiomyocytes, the effects of ISTA and LEVO were evaluated both individually and in combination to define the contribution of each intervention and their functional interaction (see supplement for more details).

### 2.1. ISTA improves myocardial contraction and Ca^2+^ cycling

#### 2.1.1. Contractile performance

In murine ventricular trabeculae, selected multicellular preparations that reflect aligned myocardial architecture, ISTA (20 µM) elicited an overall improvement in mechanical performance at 1 Hz, increasing developed force and accelerating both contraction and relaxation kinetics (**Figure 1a-e**). In addition, ISTA accelerated “mechanical restitution”, i.e. it boosted recovery of contractile force following premature stimulation (**Figure 1f**). On the other hand, ISTA did not modify force generation or myosin ATPase activity in chemically skinned preparations (**Figure 1h-o**), indicating that the agent does not directly affect sarcomeric function under these conditions. To identify the cellular mechanisms underlying these functional effects, intracellular Ca^2+^ dynamics were subsequently investigated in isolated ventricular cardiomyocytes.

**Figure 1.**
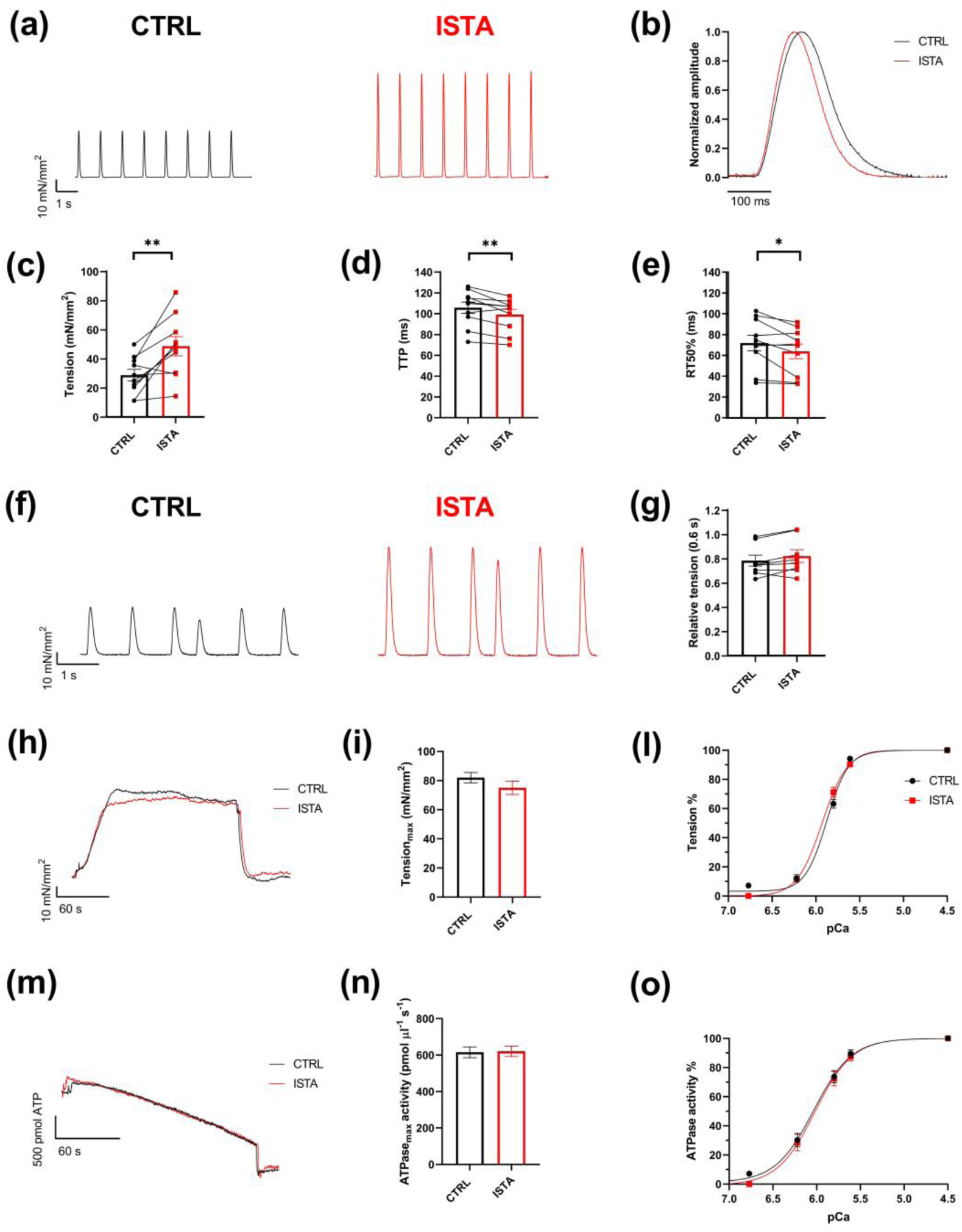
ISTA effect on contractile performance. (a) Representative force twitches from a trabecula paced at 1 Hz, at baseline and in the presence of ISTA (20 µM). (b) Normalized superimposed twitches recorded at 1 Hz, in the absence and presence of ISTA. (c) Peak isometric tension (Basal: 28.6 ± 4.1; ISTA: 48.4 ± 6.6; n=10, N=8). (d) Time to peak contraction, TTP (Basal: 105.6 ± 5.4; ISTA: 99 ± 5, n=10, N=8). (e) Time from peak to 50% relaxation, RT50% (Basal: 72.2 ± 7.5; ISTA: 64.2 ± 7; n=10, N=8). (f) Representative traces from a trabecula at baseline and with ISTA. A premature stimulus at a time interval of 0.6 s was given in a series of constant stimulation at 1 Hz. (g) Relative tension of extrasystole (at 0.6 s) normalized to steady-state twitch amplitude (Basal: 0.78 ± 0.04; ISTA: 0.82 ± 0.05; n=8, N=6). (h) Representative contraction-relaxation cycle in skinned muscle strips maximally activated in pCa 4.5. (i) Mean values of maximal tension from skinned muscle strips in the absence (n=13, N=6) and in the presence (n=16, N=3) of ISTA. (l) Force-pCa relationship in the absence (n=13, N=6) and in the presence (n=16, N=3) of ISTA (m) Representative decline of NADH absorbance as a readout of ATP consumption in skinned muscle strips maximally activated in pCa 4.5. (n) Mean values of maximal ATPase from skinned muscle strips in the absence (n=13, N=6) and in the presence (n=16, N=3) of ISTA. (o) ATPase-pCa relationship in the absence (n=13, N=6) and in the presence (n=16, N=3) of ISTA. Data are expressed as mean ± SEM. Individual data points are shown. Paired Student’s t-test, *<0.05, **<0.01.

#### 2.1.2. Ca^2+^ dynamics

ISTA (20 µM) effect on intracellular Ca^2+^ dynamics was assessed in intact CMs, firstly during steady-state field-stimulation at 1Hz (**Figure 2**). ISTA significantly increased Ca^2+^ transient (CaT) amplitude (+17%, p<0.05, Fig. 2C), accelerated both rise time (t_peak,_ -4%, p<0.05, **Figure 2d**) and decay rate (τ decay, -13%, p<0.001, **Figure 2e**), and left diastolic Ca^2+^ levels (CaD) unchanged (**Figure 2f**), thus indicating improved systolic Ca^2+^ dynamics without impairment of diastolic Ca^2+^ homeostasis.

**Figure 2.**
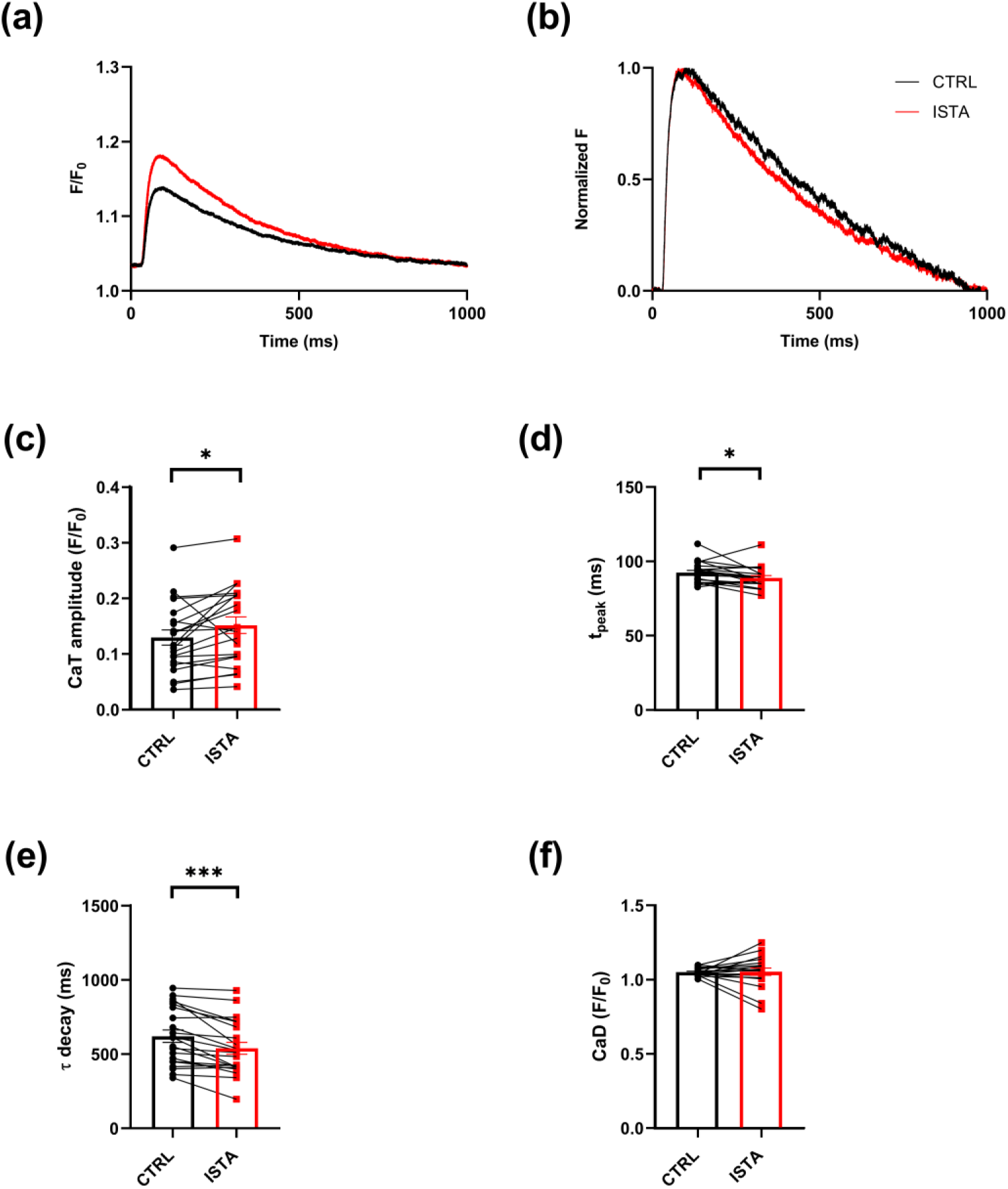
ISTA (20 µM) effect on Ca^2+^ dynamics. (a) Representative traces of Ca^2+^ transients (CaTs). (b) Normalized CaT traces emphasize changes in τ decay. (c) CaT amplitude (CTRL: 0.13 ± 0.01, n=21, N=4; ISTA: 0.15 ± 0.01, n=21, N=4). (d) t_peak_ (CTRL: 93 ± 1.53, n=21, N=4; ISTA: 89 ± 1.52, n=21, N=4). (e) τ decay (CTRL: 619 ± 42, n=21, N=4; ISTA: 538 ± 40, n=21, N=4). (f) CaD (CTRL: 1.05 ± 0,01, n=19, N=4; ISTA: 1.06 ± 0.02, n=19, N=4). Data are expressed as mean ± SEM. Paired Student’s t-test (panels c, e, f) and Wilcoxon test (panel d), * p<0.05, *** p<0.001.

An SR “reloading” protocol was then used to assess ISTA effect on dynamic Ca^2+^ compartmentalization in the SR (see methods). ISTA increased the rate of SR Ca^2+^ accumulation after caffeine-induced depletion (p<0.05, **Figure S1c, d**). Furthermore, ISTA decreased τ decay (p<0.001, **Figure S1e**) and t_peak_ (p<0.0001, **Figure S1g**), with the reduction in t_peak_ being most pronounced during the initial loading steps. Together, these findings are consistent with more efficient SERCA2a-mediated Ca^2+^ reuptake and retention within the SR. In apparent contrast with its effect on progressive SR reloading, during steady-state stimulation ISTA changed neither SR Ca^2+^ content (CaSR) nor fractional release (**Figure S1l, m**). NCX-dependent Ca^2+^ extrusion (**Figure S2**) and I_CaL_ restitution properties (**Figure S3**) were also unaffected, thus assigning its effect on Ca^2+^ cycling to improved SR function rather than altered sarcolemmal Ca^2+^ transport.

Collectively, these findings demonstrate that 20 µM ISTA generates a coherent functional phenotype characterized by enhanced SR Ca^2+^ cycling and improved mechanical performance. This phenotype was therefore used as the reference condition to determine whether the subsequent addition of LEVO could provide further functional benefit.

### 2.2 LEVO fails to provide additional benefit over the ISTA-induced phenotype

#### 2.2.1. Contractile performance

LEVO (0.5 µM) was added on top of ISTA (20 µM) in twitching muscles. Unexpectedly under these conditions, LEVO did not produce any significant further increase in developed force at 1 Hz, and in the large majority of the muscles (7/8), LEVO caused a modest (> 5%), although significant, drop in contraction amplitude (**Figure 3a-c**). Contraction kinetics were essentially unaffected and remained comparable before and after the addition of LEVO (**Figure 3d, e**). Likewise, the improvement in mechanical restitution induced by ISTA was not significantly enhanced by the subsequent addition of LEVO (**Figure 3f, g**).

**Figure 3.**
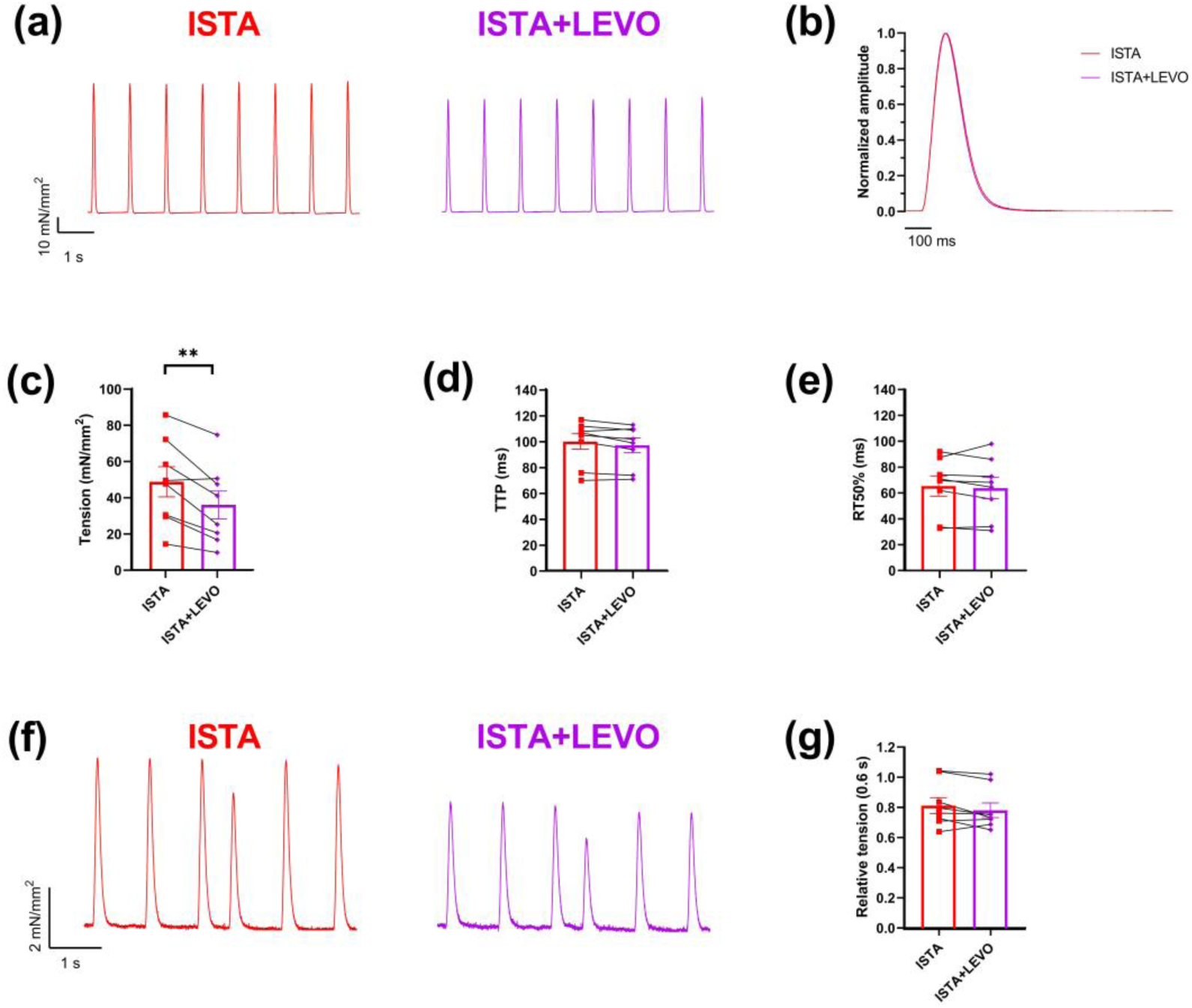
Combined ISTA+LEVO effect on contractile performance. (a) Representative force twitches from a trabecula paced at 1 Hz, in the presence of ISTA and ISTA+LEVO. (b) Normalized superimposed twitches recorded at 1 Hz, with ISTA and ISTA+LEVO. (c) Peak isometric tension (ISTA: 48.5 ± 8.3; ISTA+LEVO: 35.8 ± 7.6; n=8, N=6). (d) Time to peak contraction, TTP (ISTA: 99.4 ± 6; ISTA+LEVO: 96.5 ± 5.7, n=8, N=6). (e) Time from peak to 50% relaxation, RT50% (ISTA: 65.2 ± 7.8; ISTA+LEVO: 63.7 ± 8.2; n=8, N=6). (f) Representative traces from a trabecula with ISTA and with ISTA+LEVO. A premature stimulus at a time interval of 0.6 s was given in a series of constant stimulation at 1 Hz. (g) Relative tension of extrasystole (at 0.6 s) normalized to steady-state twitch amplitude (ISTA: 0.82 ± 0.05; ISTA+LEVO: 0.79 ± 0.05; n=8, N=6). Data are expressed as mean ± SEM. Individual data points are shown. Paired Student’s t-test, **<0.01.

#### 2.2.2. Ca^2+^ dynamics

In a separate group of cardiomyocytes, the effects of simultaneous ISTA+LEVO application were assessed against the corresponding drug-free controls (**Figure 4**). Under steady-state stimulation at 1 Hz ISTA+LEVO significantly reduced τ decay (-16%, p<0.01, **Figure 4e**) and CaD (-7%, p<0.05, **Figure 4f**), without changing CaT amplitude and t_peak_ (**Figure 4c, d**). SR Ca²⁺ content and reuptake rate, were not affected by the combination of the two drugs (**Figure S4**). NCX-dependent Ca²⁺ extrusion (**Figure S5**) and I_CaL_ restitution (**Figure S6**) also remained unaltered, indicating that the lack of additional mechanical benefit was paralleled by the absence of measurable enhancement of intracellular Ca^2+^ cycling.

**Figure 4.**
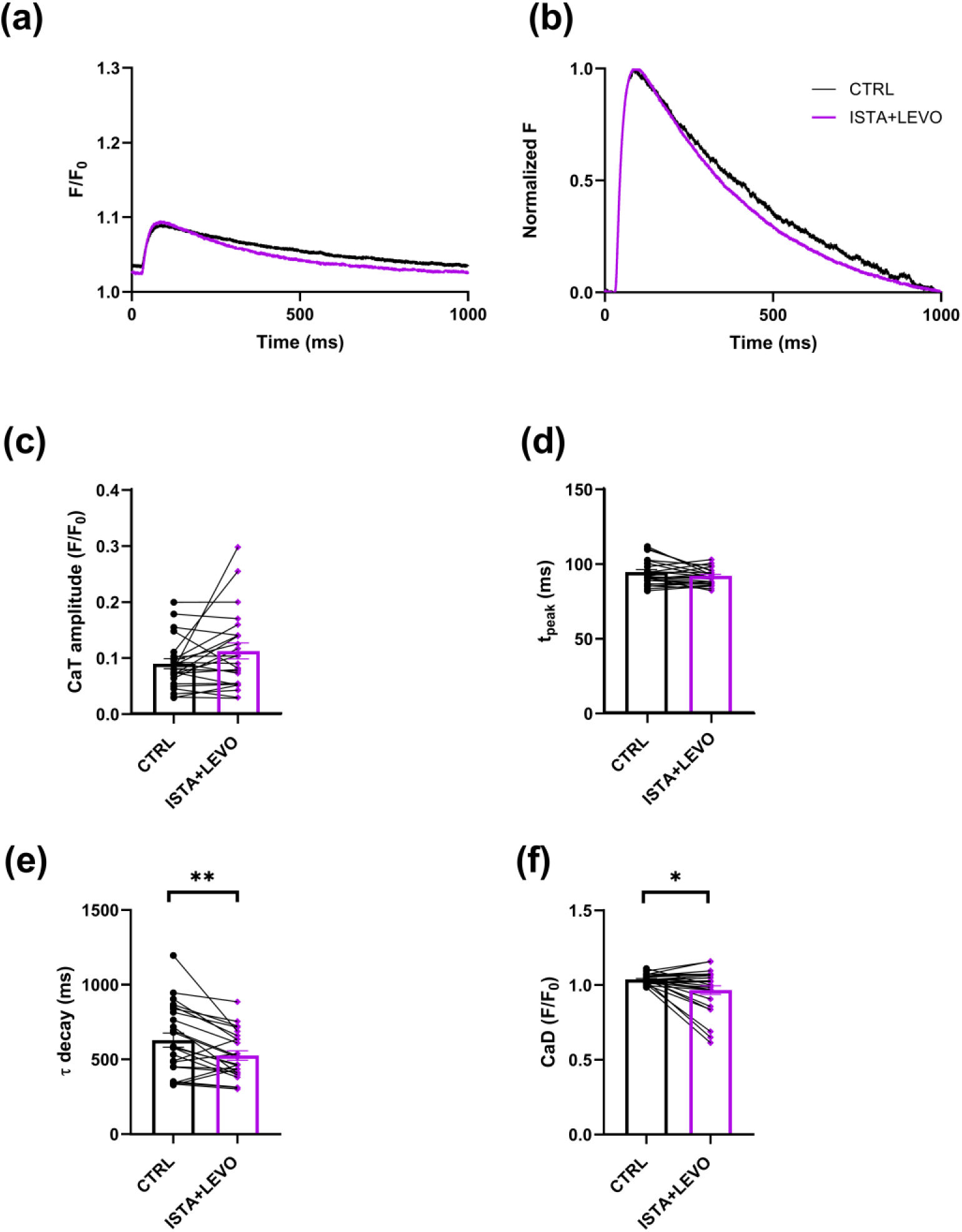
ISTA+LEVO effect on Ca^2+^ dynamics. (a) Representative traces of CaTs. (b) Normalized CaT traces. (c) CaT amplitude (CTRL: 0.088 ± 0.01, n=24, N=6; ISTA+LEVO: 0.11 ± 0.01, n=24, N=6). (d) t_peak_ (CTRL: 93 ± 1.66, n=25, N=6; ISTA+LEVO: 91 ± 1.07, n=25, N=6). (e) τ decay (CTRL: 630 ± 47, n=24, N=6; ISTA+LEVO: 527 ± 32, n=24, N=6). (f) CaD (CTRL: 1.04 ± 0.01, n=25, N=6; ISTA+LEVO: 0.97 ± 0.03, n=25, N=6). Data are expressed as mean ± SEM. Paired Student’s t-test (panels d, e) and Wilcoxon test (panel c, f), * p<0.05, ** p<0.01.

Overall, adding 0.5 µM LEVO to ISTA-treated trabeculae provided no further mechanical benefit, while simultaneous ISTA+LEVO application in cardiomyocytes accelerated Ca^2+^ transient decay and reduced diastolic Ca^2+^ relative to controls. To determine whether the additional mechanical benefit could reflect limited intrinsic responsiveness to LEVO, its effects were subsequently examined in the absence of ISTA using both multicellular ventricular preparations and isolated cardiomyocytes.

### 2.3 Inconsistent efficacy of LEVO in murine myocardium

#### 2.3.1. Contractile performance

LEVO at the selected human therapeutic dose of 0.5 µM elicited a markedly heterogeneous mechanical response in twitching mouse trabeculae at 1 Hz. The majority of the multicellular preparations (3/5) exhibited a clear positive inotropic effect, whereas the remaining preparations showed a reduction in developed force (**Figure 5a, c**). As a result, the treatment produced no overall effect on contraction amplitude. Similarly, twitch contraction kinetics were not affected by LEVO (**Figure 5b, d, e**). Mechanical restitution was studied in a subset of trabeculae: LEVO did not cause any change in twitch recovery with respect to baseline (**Figure 5f**). Importantly and unexpectedly, Force-PCa curves obtained in skinned muscle strips show no variations in myofilament Ca^2+^ sensitivity with LEVO, indicating that the drug, at this dose, does not act as a Ca^2+^ sensitizer in murine myocardium (**Figure 5h**).

**Figure 5.**
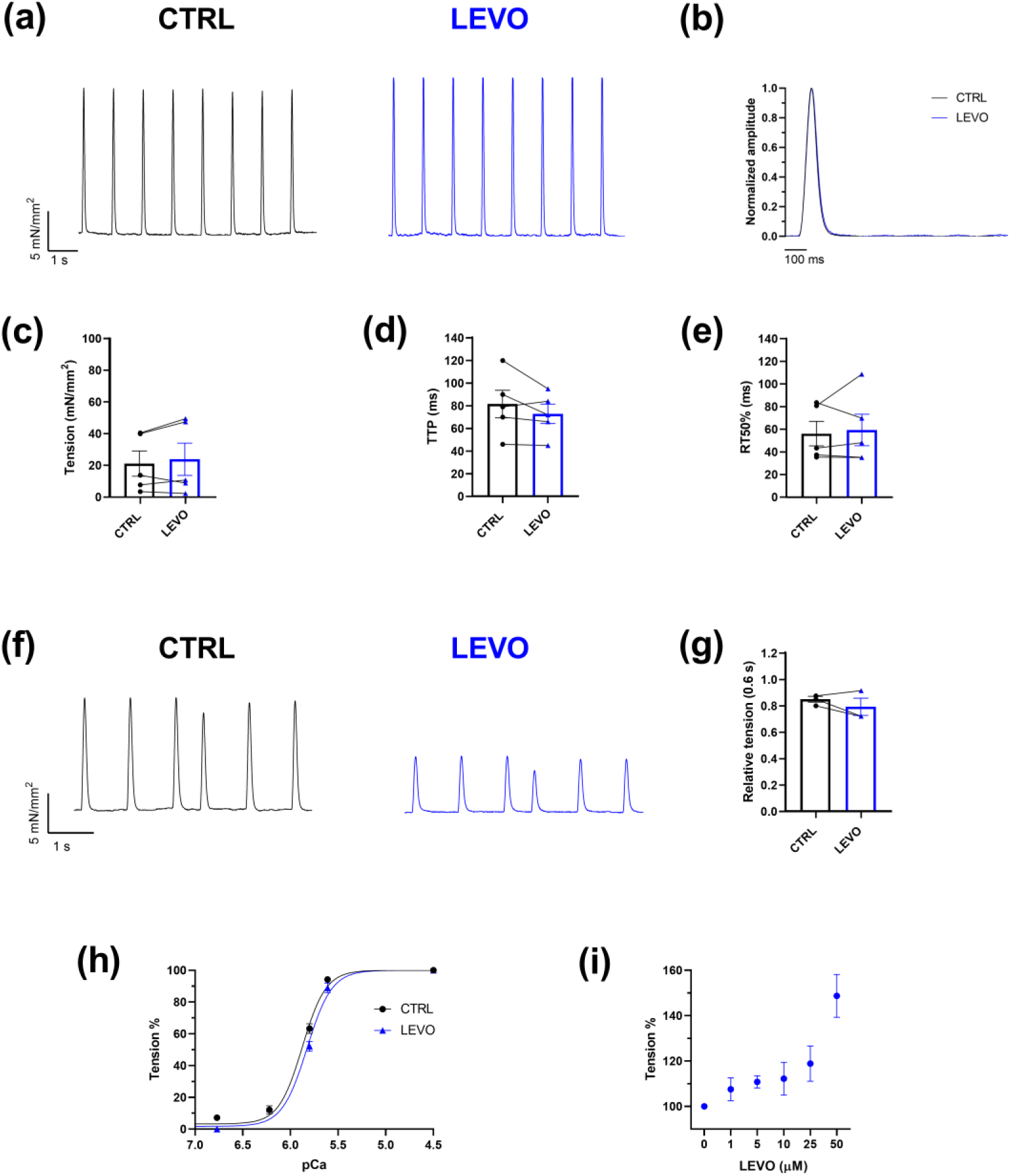
LEVO effect on contractile performance. (a) Representative force twitches from a trabecula paced at 1 Hz, at baseline and in the presence of LEVO. (b) Normalized superimposed twitches recorded at 1 Hz, in the absence and presence of LEVO. (c) Peak isometric tension (Basal: 21.1 ± 8; LEVO: 23.8 ± 10.2; n=5, N=3). (d) Time to peak contraction, TTP (Basal: 81 ± 12.1; LEVO: 72.4 ± 8.5, n=5, N=3). (e) Time from peak to 50% relaxation, RT50% (Basal: 56.1 ± 10.7; LEVO: 59.3 ± 13.9; n=5, N=3). (f) Representative traces from a trabecula at baseline and with LEVO. A premature stimulus at a time interval of 0.6 s was given in a series of constant stimulation at 1 Hz. (g) Relative tension of extrasystole (at 0.6 s) normalized to steady-state twitch amplitude (Basal: 0.84 ± 0.02; LEVO: 0.79 ± 0.06; n=3, N=2). (h) Tension–pCa relationships in skinned muscle strips under CTRL conditions (n=13, N=3) and in the presence of LEVO (n=5, N=1). Tension is expressed as a percentage of maximal Ca^2+^-activated tension. (i) Concentration–response relationship for LEVO in skinned muscle strips activated at a submaximal Ca^2+^ concentration (pCa 5.8), with tension expressed as a percentage of the corresponding drug-free value (n=16, N=3). Data are expressed as mean ± SEM.

To further investigate whether LEVO induced myofilament sensitization to Ca^2+^, as postulated by its class action, cell shortening (twitch) and cytosolic Ca^2+^ were simultaneously measured in isolated patch-clamped ventricular myocytes. Sets of voltage steps, to variably activate I_CaL_, were applied to elicit Ca^2+^ transients and twitches of variable amplitudes. Plots of twitch vs CaT amplitude were thus obtained to assess myofilament response to Ca^2+^. Consistent with the heterogeneous response observed in multicellular ventricular preparations, LEVO induced a clear cut leftward shift of the twitch/CaT plot (increased Ca^2+^ sensitivity) in only a minority (25%, n=2/8) of cardiomyocytes (**Figure S7**).

For completeness, sarcomere modulation by LEVO was also tested over a broad range of concentrations in a subset of skinned multicellular preparations, activated at sub-maximal level (pCa 5.8). An increase of developed tension was indeed observed, but only at LEVO concentrations exceeding therapeutically achievable ones by three orders of magnitude (**Figure 5i**).

#### 2.3.2. Ca^2+^ dynamics

We next assessed the effects of LEVO on intracellular Ca²⁺ dynamics during steady-state stimulation at 1 Hz (**Figure 6**). LEVO did not significantly affect CaT amplitude, t_peak_ or diastolic Ca^2+^ levels (**Figure 6c, d, f**), but unexpectedly accelerated Ca^2+^ transient decay (τ decay -13%, p<0.01, **Figure 6e**), an effect like that exerted by ISTA and compatible with SERCA2a stimulation; however, in the SR reloading protocol LEVO failed to significantly accelerate SR reloading (**Figure S8c, d**). SR Ca^2+^ content, NCX-dependent Ca^2+^ extrusion (**Figure S9**) and I_CaL_ restitution (**Figure S10**) were unaffected by LEVO.

**Figure 6.**
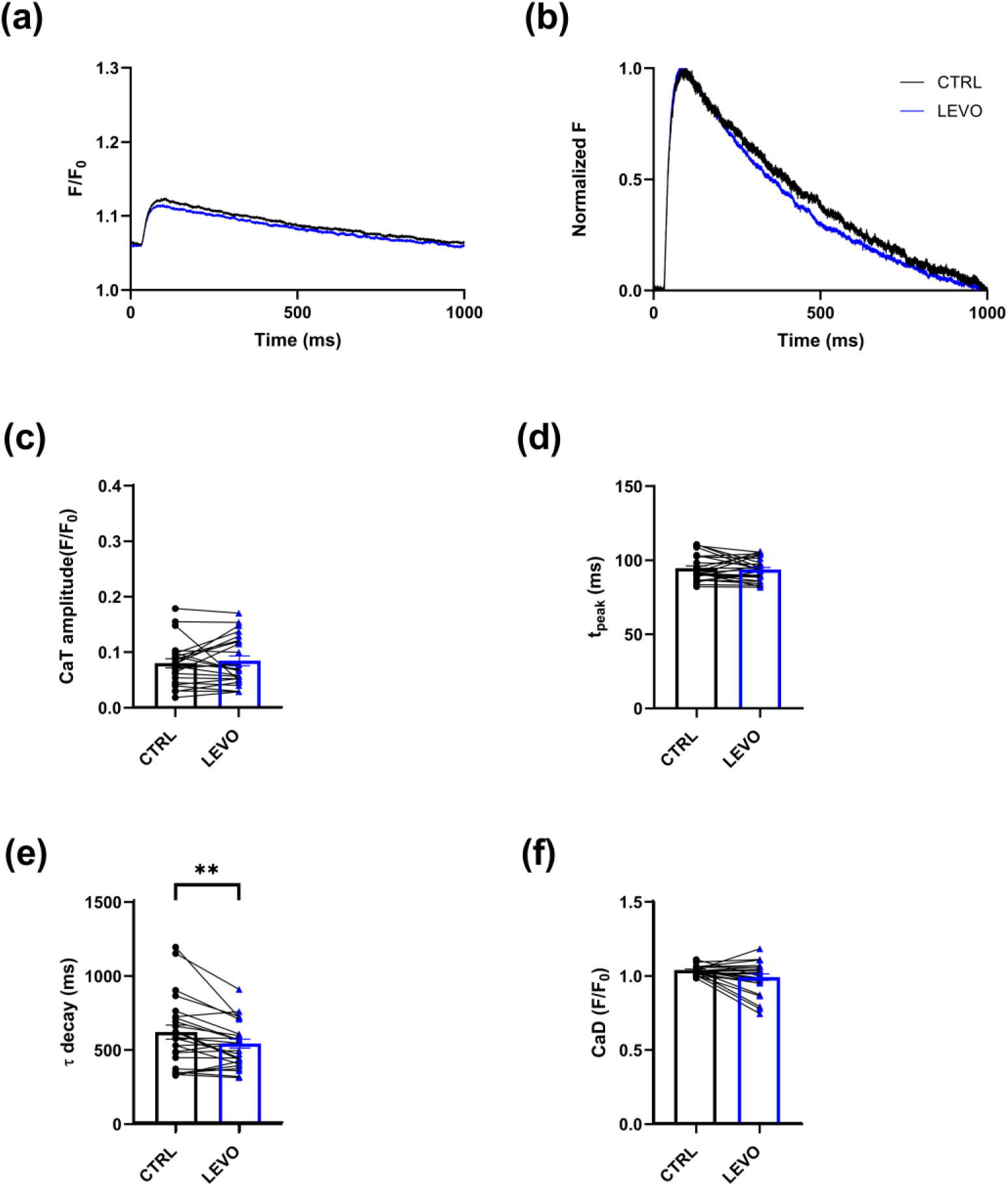
LEVO (0.5 µM) effect on Ca^2+^ dynamics. (a) Representative traces of Ca^2+^ transients (CaT). (b) Normalized CaT traces. (c) CaT amplitude (CTRL: 0.08 ± 0.01, n=24, N=4; LEVO: 0.08 ± 0.01, n=24, N=4). (d) t_peak_ (CTRL: 94 ± 1.70, n=26, N=4; LEVO: 94 ± 1.47, n=26, N=4). (e) τ decay (CTRL: 609 ± 47, n=25, N=4; LEVO: 532 ± 29, n=25, N=4). (f) CaD (CTRL: 1.03 ± 0,01, n=24, n=4; LEVO: 0.98 ± 0.022, n=24, N=4). Data are expressed as mean ± SEM. Paired Student’s t-test (panels c, d, e) and Wilcoxon test (panel f), ** p<0.01.

While acceleration of Ca^2+^ transient decay alone would hardly support positive inotropy, the question arose whether it could be a less apparent consequence of myofilament modulation by LEVO (e.g. increased Ca^2+^ buffering) or rather reflect an ancillary action. LEVO ancillary properties include inhibition of phosphodiesterase-3 (PDE3) [16,17]; hence, we tested the hypothesis that the latter might account for the acceleration of CaT decay.

#### 2.3.3. Contribution of PDE3 inhibition to LEVO effect on Ca^2+^ dynamics

Because LEVO is known to possess phosphodiesterase-3 (PDE3) inhibitory activity we next investigated whether the acceleration of τ decay induced by LEVO could be attributed to PDE3 inhibition. To this end, the effects of LEVO were compared with those of the selective PDE3 inhibitor cilostamide (CILO, 0.5 µM) [17], and the ability of LEVO to elicit additional functional responses following PDE3 inhibition was evaluated.

Like LEVO, CILO reduced τ decay (-18%, p<0.001, **Figure 7e**) without significantly affecting other Ca^2+^ transient parameters. Notably, in the presence of CILO, LEVO failed to further modulate Ca^2+^ dynamics (**Figure 7**). These results suggest that the acceleration of τ decay induced by LEVO was largely mediated by its PDE3 inhibitory activity.

**Figure 7.**
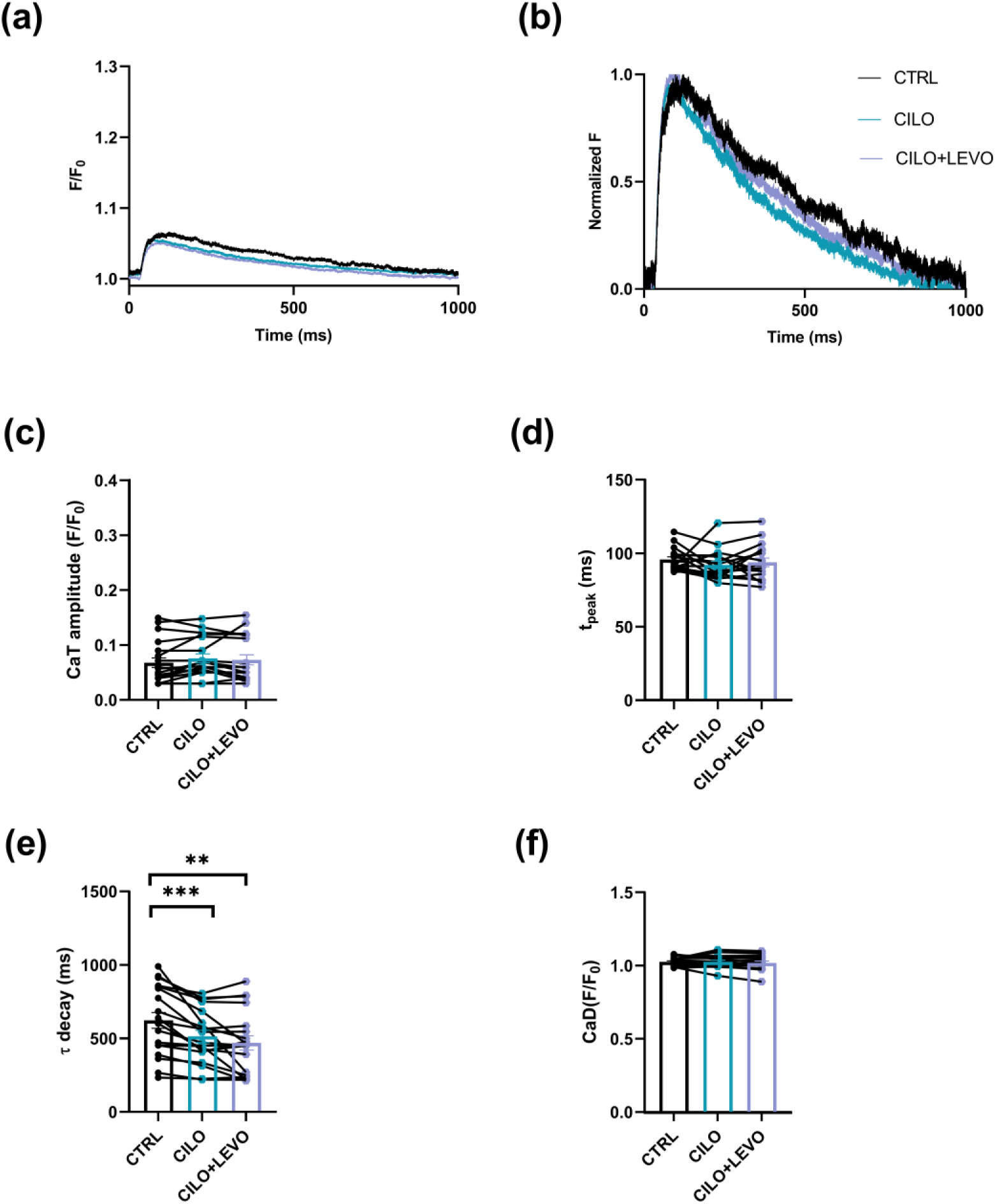
CILO (0.5 µM) and CILO+LEVO effect on Ca^2+^ dynamics. (a) Representative traces of Ca^2+^ transients (CaT). (b) Normalized CaT traces. (c) CaT amplitude (CTRL: 0.07 ± 0.009, n=19; CILO: 0.08 ± 0.008, n=19, N=2; CILO+LEVO: 0.07 ± 0.009, n=19, N=2). (d) t_peak_ (CTRL: 96 ± 1.88, n=17, N=2; CILO: 92 ± 2.49, n=17, N=2; CILO+LEVO: 94 ± 2.97, n=17, N=2). (e) τ decay (CTRL: 625 ± 54, n=19, N=2; CILO: 515 ± 42, n=19, N=2; CILO+LEVO: 471 ± 49, n=19, N=2). (f) CaD (CTRL: 1.03 ± 0.007, n=16, N=2; CILO: 1.03 ± 0.011, n=16, N=2; CILO+LEVO: 1.02 ± 0.013, N=16, N=2). Data are expressed as mean ± SEM. One-way ANOVA followed by Tukey’s post-hoc test, ** p<0.01, *** p<0.001.

## 3. Discussion

The present study was designed to elucidate the mechanisms underlying the positive inotropic interaction between ISTA and LEVO recently observed in pilot experiments conducted on human failing myocardium. Unexpectedly, this objective could only be partially achieved because therapeutic LEVO concentrations failed to elicit the anticipated positive inotropic response in murine preparations. Nevertheless, the study provided insight into the effects of the two agents on myocardial function and Ca^2+^ handling, while highlighting limitations of murine preparations for investigating their functional interaction.

### Responsiveness to LEVO in murine myocardium

LEVO improves cardiac function in patients with acute and advanced heart failure [18] and, in our own hands, it enhanced force development in ex-vivo multicellular human myocardial preparations. This observation is consistent with previous findings in myocardial preparations from humans and other mammalian species expressing ventricular β-myosin, such as guinea pigs, in which LEVO (0.03–10 μmol/L) reversibly increased force at submaximal activating free Ca^2+^ concentrations without significantly affecting maximal force [19]. To our knowledge, the present study provides the first direct evidence that murine myocardium displays an erratic contractile responsiveness to LEVO at clinically relevant concentrations (**Figure 5**); because the effect mediated by PDE3 inhibition (acceleration of CaT decay) was consistently observed instead, lack of inotropy was likely the consequence of lack of sarcomere sensitization. This suggests that the latter may be essential in accounting for LEVO-induced inotropy, thus contributing to the long-standing debate of whether PDE3 inhibition may prevail instead [17,20].

Notably, a relevant increase in sarcomere sensitivity to Ca^2+^ (+ 20-50% of baseline tension) was observed in skinned murine ventricular preparations when LEVO was tested at concentration substantially higher than clinically relevant ones (**Figure 5i**). While this finding indicates that murine sarcomeres are capable of responding to LEVO, albeit only at supra-therapeutic concentrations.

### Potential mechanisms underlying the reduced murine responsiveness

The reduced responsiveness of murine myocardium to LEVO is species-specific and likely multifactorial. LEVO enhances contractility by stabilizing the Ca^2+^-bound conformation of cardiac troponin C, thereby increasing myofilament Ca^2+^ sensitivity and promoting force generation without increasing the amplitude of the intracellular Ca^2+^ transient [21,22]. The efficacy of this mechanism depends on the characteristics of the contractile apparatus and on the kinetics of intracellular Ca^2+^ cycling. Compared with human myocardium, murine cardiomyocytes exhibit shorter systolic activation [23], supported by markedly faster contraction-relaxation kinetics, largely dependent on sarcoplasmic reticulum Ca^2+^ uptake through SERCA2a. These characteristics may reduce the temporal window during which stabilization of the Ca^2+^-troponin C complex can effectively augment force generation.

In addition, species-dependent differences in sarcomeric protein composition (e.g. prevalence in the mouse of α versus β myosin isoform) and post-translational regulation (e.g. high phosphorylation levels of myofilament proteins) [24] may further influence the responsiveness of the contractile apparatus to Ca^2+^ sensitization. Collectively, these features may reduce the functional contribution of myofilament Ca^2+^ sensitization in mice. Although this interpretation remains speculative, it provides a plausible explanation for reconciling the limited effects observed in murine myocardium with the well-established efficacy of LEVO in human cardiac tissue or rodents expressing β myosin (e.g. guinea pigs) [19].

### Cellular heterogeneity of the LEVO response

A variable contractile response to LEVO might arise from differences at the myocyte level or in tissue architecture (e.g. affecting drug diffusion under tissue perfusion). In the present study LEVO contractile responsiveness was similarly heterogeneous between multicellular preparations (**Figure 5**) and isolated myocytes (**Figure S7**), suggesting that tissue-level factors alone cannot explain this variability. Although its cellular determinants remain unresolved, this heterogeneity may contribute to the lack of a consistent positive inotropic response to LEVO under our experimental conditions.

### PDE3 inhibition accounts for the effects of LEVO on Ca^2+^ handling

The selective PDE3 inhibitor CILO reproduced the effect of LEVO on τ decay, and subsequent administration of LEVO failed to induce any further acceleration (**Figure 7**). This assigns LEVO effect on τ decay to PKA-dependent relief of PLN-induced SERCA2a restraint [25]. This finding also implies that the PDE3-inhibitory action of LEVO, albeit not associated with a measurable inotropic response, was present in murine myocytes under our experimental conditions. Whereas PKA activation modulates Ca^2+^ dynamics also through targets other than SERCA2a (CaV1.2, RyRs etc.), acceleration of CaT decay was the only significant LEVO effect on Ca^2+^ transients. This might reflect the selective expression of the PDE3 isoform in the subcellular compartment relevant to SERCA2a modulation [26].

### Implications for the interaction between ISTA and LEVO

These observations also provide a possible explanation for the absence of an additive interaction between ISTA and LEVO. ISTA positive inotropic effect, clearly observed also in multicellular preparations, was associated with changes in Ca^2+^ dynamics (acceleration of τ decay, faster SR reloading) compatible with SERCA2a stimulation. In the case of ISTA, the latter occurs through PKA-independent phospholamban (PLN) antagonism [8]. The outcome of measurements performed in the presence of PDE3 inhibition suggests that acceleration of τ decay by LEVO may also reflect SERCA2a stimulation through relief of PLN inhibitory effect, albeit achieved through PKA signaling instead [27]. Irrespective of the primary mechanism, ISTA and LEVO may converge on the same target in modulating Ca^2+^ dynamics, which may make their action mutually exclusive.

Assuming that subcellular expression of the PDE3 isoform holds true for human myocytes [28], the summation of ISTA and LEVO inotropic effects preliminary observed in human preparations more likely relies on LEVO-induced sarcomere sensitization to Ca^2+^, rather than on PDE3 inhibition. PKA-mediated TnI phosphorylation notoriously reduces sarcomere affinity for Ca^2+^ [29]; therefore, if anything, PDE inhibition would antagonize sarcomere sensitization to Ca^2+^.

In practical terms, the present findings suggest that the mouse, in which the PDE3 inhibitory effect is the only present at therapeutic concentration, may not be a suitable model for investigating the Ca^2+^-sensitizing properties of LEVO.

## 4. Conclusions

While unsuitable to analyze the mechanism underlying the additive functional interaction, previously observed between ISTA and LEVO in human failing myocardium, the present study unveils markedly reduced responsiveness of mouse myocardium to LEVO-induced sarcomere sensitization to Ca^2+^. Rather than challenging the concept of ISTA-LEVO complementarity in inotropic therapy, these findings identify an important species-dependent limitation of the murine model in investigating LEVO-induced inotropism. The results also indicate that the PDE3 inhibitory effect of LEVO is retained in murine myocytes, without resulting in significant inotropy. This may stress the role of sarcomere sensitization in human myocardium, in which LEVO does increase contractility and positively interacts in this effect with ISTA.

## 5. Materials and Methods

### 5.1 Experimental model

The experiments were performed on cardiomyocytes (CMs) or multicellular preparations isolated from wild-type mice aged 8 to 12 weeks. CMs were enzymatically dissociated with a manual perfusion method [30] which does not discriminate between right and left ventricles (see supplement for details) .

### 5.2 Experimental techniques and protocols

#### Isolated CMs

- Cytosolic Ca^2+^ was optically measured by using Fluo 4-AM (10 µM) as the Ca^2+^-sensitive probe; fluorescence (F) was normalized to the value measured after prolonged quiescence (F_0_). Ca^2+^measurements were carried out in field-stimulated or voltage-clamped CMs, as specified in the text. The following parameters were measured in field-stimulated CMs during steady-state stimulation at 1 Hz: Ca^2+^ transient (CaT) amplitude, CaT decay kinetics (τ decay), CaT rise-time (t_peak_) and diastolic Ca^2+^ (CaD).
- The Ca^2+^ uptake function of the sarcoplasmic reticulum (SR) was assessed using an “SR reloading” protocol performed under voltage-clamp conditions, as previously described [14,31]. Briefly, the SR Ca^2+^ stores were first depleted by a caffeine (10 mM) pulse and progressively refilled through a train of voltage steps (-40/0 mV at 0.25 Hz). CaTs and voltage-gated L-type calcium current (I_CaL_) were simultaneously recorded at each step of the protocol and the following parameters were quantified: i) CaT amplitude, ii) CICR gain (defined as the ratio between SR Ca^2+^ release and Ca²⁺ influx through ICaL), iii) τ decay, iv) t_peak_, and v) CaD.
- SR Ca^2+^ content (CaSR) was estimated in voltage-clamped CMs from the integral of the membrane current (mostly representing I_NCX_) elicited by the caffeine pulse, applied after a loading train of voltage steps (−40/0 mV at 1 Hz).
- The fraction of SR Ca²⁺ released following membrane depolarization (fractional release) was quantified as the ratio between the amplitudes of voltage-evoked and caffeine-induced CaTs.
- NCX functional properties were derived from linear regression analysis of the trailing portion of the I_NCX_/[Ca^2+^] loops generated during caffeine-induced CaTs. The slope coefficient and the 0 I_NCX_ intercept were considered surrogate measures of NCX “conductance” and cytosolic [Ca²⁺] at electrochemical equilibrium, respectively.
- I_CaL_ restitution was evaluated in voltage-clamped CMs by applying a protocol consisting of a conditioning stimulation at 1 Hz (s_1_) followed by an extra-stimulus (s_2_). The s_1_-s_2_ interval was progressively increased from 10 to 800 ms. The ratio between the current elicited by extra-stimulus and that obtained during the steady-state stimulation (s_2_/s_1_) was plotted as a function of the s_1_-s_2_ interval to obtain the restitution curves of I_CaL._
- Ca^2+^ sensitivity of contraction was determined in voltage-clamped CMs by applying depolarizing voltage steps of progressively increasing amplitude (V_hold_ = -40 mV) to elicit graded CaTs and twitch contractions. The twitch/CaT relationship was obtained by normalizing both twitch amplitude and CaT amplitude to the respective steady-state values recorded under CTRL conditions.

#### Multicellular preparations

- Twitch contraction and sarcomere mechanics and energetics were assessed in intact trabeculae and in skinned muscle strips, respectively, as previously described [32,33].
- Right ventricular trabeculae were carefully dissected from explanted mouse hearts. Then they were mounted between a basket-shaped end of a force transducer and a motor, controlled by custom-made software (LabVIEW).
- Intact muscles were gradually stretched to the optimal length and allowed to stabilize for 25-30 minutes at 37°C, 1.5 mmol/L [CaCl_2_], and a stimulation frequency of 1 Hz (baseline conditions).
- Isometric force and kinetics (time to peak contraction, TTP, and time from peak to 50% relaxation, RT50) were evaluated at baseline conditions and following the administration of Istaroxime 20 µM (ISTA), Levosimendan 0.5 µM (LEVO) or their combination (ISTA+LEVO).
- Mechanical restitution was studied by giving a premature stimulus at different time intervals (25-800 ms) in a series of constant stimulation at 1 Hz, to elicit a premature contraction (extrasystole). The amplitude of extrasystole was reduced and progressively increased with increasing time intervals, until the steady-state level was again reached. The relative amplitudes of extrasystoles were plotted against the corresponding stimulus intervals, and a single-exponential fit was applied to the restitution curve to estimate the restitution rate.
- Skinned trabeculae were obtained by overnight incubation (4°C) in a relaxing solution (pCa 9) containing 0.5% Triton X100. After Triton removal, a selected skinned trabecula was then attached to aluminum T-clips and mounted in the experimental setup. Before the first activation-relaxation cycle, the sarcomere length of the preparation was adjusted to 2.2 µm.
- Skinned trabeculae were exposed to solutions containing different Ca²⁺ concentrations at 21 °C to evaluate the pCa-force relationship. Sarcomere energetics was assessed by simultaneous measurement of isometric force and ATP consumption using an enzyme-coupled assay [34].

### 5.3 Statistical analysis

Statistical analysis was carried out with GraphPad Prism 8. Normality of distribution was assessed using D’Agostino-Pearson’s normality test. Comparison of two sample means was carried out with paired parametric (Student’s t) or non-parametric (Wilcoxon) tests, according to the data distribution. Comparisons among three conditions were carried out with one-way ANOVA (Tukey’s post-hoc correction). In the case of repeated measurements (SR loading and restitution protocols), a mixed-effects model (Sidak post-hoc correction) containing “Treatment” (CTRL vs drug) and “Variable” (step # or s1-s2 interval) factors was used. Significance of “Treatment X Variable interaction” (i.e. difference between Treatments in their response to the Variable) was tested. Data are presented as mean ± SEM. For paired data, lines connect pre- and post-treatment measurements obtained from the same cell or muscle preparation. For each experiment, the number of cells or preparations (n) and the number of animals from which they were obtained (N) are indicated in the respective figure legend. P values <0.05 were considered statistically significant.

## Supplementary Materials

The following supporting information can be downloaded at https://www.mdpi.com/article/doi/s1, Supplementary Methods; Figure S1: ISTA (20 μM) effect on SR Ca^2+^ content and uptake; Figure S2: ISTA effect on the relationship between I_NCX_ and [Ca^2+^]; Figure S3: ISTA effect on I_CaL_ restitution; Figure S4: ISTA+LEVO effect on SR Ca^2+^ content and uptake; Figure S5: ISTA+LEVO effect on the relationship between I_NCX_ and [Ca^2+^]; Figure S6: ISTA+LEVO effect on I_CaL_ restitution; Figure S7: Effects of LEVO on contractile performance in isolated cardiomyocytes; Figure S8: LEVO effect on SR Ca^2+^ content and uptake; Figure S9: LEVO effect on the relationship between I_NCX_ and [Ca^2+^]; Figure S10: LEVO effect on I_CaL_ restitution.

## Author Contributions

Conceptualization, C.P., A.Z., G.M., C.F. and F.L.; methodology, C.P., J.M.P., A.Z., C.F. and F.L.; investigation, F.M., N.G., C.M., M.D., M.S. and J.M.P.; formal analysis, F.M., N.G., C.M., M.D. and M.S.; validation, F.M., N.G., C.M., M.D., M.S., C.F. and F.L.; data curation, F.M., N.G., C.M., M.D., M.S., C.F. and F.L.; visualization, F.M., N.G., C.M., M.D. and M.S.; resources, A.Z., C.F. and F.L.; writing—original draft preparation, F.L.; writing—review and editing, all authors; supervision, A.Z., C.F. and F.L.; project administration, C.F. and F.L.; funding acquisition, C.F. and F.L. All authors have read and agreed to the published version of the manuscript. F.M., N.G. and C.M. contributed equally to the experimental work.

## Funding

This research was funded by the Italian Ministry of Universities and Research through the PRIN 2022 project (ID 2022-NAZ-0595) to C.F. and F.L.

## Institutional Review Board Statement

The animal study protocols were approved by the Ethics Committee for Animal Experimentation of the University of Milano-Bicocca (protocol code: 29C09.N.6AR; date of approval: 28/06/2023) and of the University of Florence (protocol code: 0DD9B.N.BXP; date of approval: 18/10/2024.

## Data Availability Statement

The datasets used and/or analyzed during the current study are available from the corresponding author upon reasonable request.

## Conflicts of Interest

The authors declare no conflicts of interest.

## Abbreviations

The following abbreviations are used in this manuscript:

ANOVA: Analysis of variance
ATP: Adenosine triphosphate
ATPase: Adenosine triphosphatase
CaD: Diastolic Ca^2+^ level
Ca_SR_: Sarcoplasmic reticulum Ca^2+^ content
CaT: Ca^2+^ transient
Ca_V_1.2: Voltage-gated L-type Ca^2+^ channel subtype 1.2
CICR: Ca^2+^-induced Ca^2+^ release
CILO: Cilostamide
CMs: Cardiomyocytes
CTRL: Control
EC: Excitation–contraction
HF: Heart failure
I_CaL_: L-type Ca^2+^current
I_NCX_: Na^+^/Ca^2+^ exchanger current
ISTA: Istaroxime
LEVO: Levosimendan
NADH: Reduced nicotinamide adenine dinucleotide
NCX: Na^+^/Ca^2+^ exchanger
pCa: Negative base-10 logarithm of the molar free Ca^2+^ concentration
PDE: Phosphodiesterase
PDE3: Phosphodiesterase type 3
PKA: Protein kinase A
PLN: Phospholamban
RT_50_: Time from peak contraction to 50% relaxation
RyRs: Ryanodine receptors
SEM: Standard error of the mean
SERCA2a: Sarco/endoplasmic reticulum Ca^2+^-ATPase isoform 2a
SR: Sarcoplasmic reticulum
TnI: Troponin I
t_peak_: Time to peak of the Ca^2+^ transient
TTP: Time to peak contraction
τ decay: Time constant of Ca^2+^ transient decay

## Notes

### Competing Interest Statement

The authors have declared no competing interest.

