## Supplement for "Species-Dependent Limitations of Murine Myocardium for Investigating the Combined Inotropic Effects of SERCA2a Stimulation and Myofilament Ca^2+^ Sensitization"

### SUPPLEMENTARY METHODS

#### Cardiomyocyte isolation

Mice were intraperitoneally anesthetized with mixture of 20 mg of ketamine-xylazine (130–7.5 mg/kg i.p), 200 µL of Seleparin (Nadroparin Calcium 9500 U.I, ItalFarmaco) and 200 µL of saline (0.9% NaCl 1M in MilliQ), euthanized by cervical dislocation. Ventricular cardiomyocytes (CMs) were isolated using a manual perfusion technique [1]. Briefly, the right ventricle was perfused with an EDTA-based buffer (130 mM NaCl, 5 mM KCl, 0.5 mM $NaH_2PO_4$, 10 mM HEPES, 10 mM Glucose, 10 mM 2,3-Butanedione Monossime, BDM, 10 mM Taurine, 5 mM EDTA; pH=7.8 with NaOH), after which the aorta was clamped and the heart removed. Enzymatic digestion was performed by sequential perfusion of the left ventricle with EDTA buffer, perfusion buffer (130 mM NaCl, 5 mM KCl, 0.5 mM $NaH_2PO_4$, 10 mM HEPES, 10 mM Glucose, 10 mM 2,3-Butanedione Monossime, BDM, 10 mM Taurine, 1 mM $MgCl_2$; pH=7.8 with NaOH), and a Liberase/trypsin solution (0.1 mg/ml Liberase + 0.02 mg/ml Trypsin EDTA and 6.25 µM $CaCl_2$). Cells were then gently dissociated, and enzymatic activity was stopped with a serum-containing buffer (10% FBS). The cell suspension was filtered, centrifuged, resuspended in $Ca^{2+}$-free solution (130 mM NaCl, 5.4 mM KCl, 0.4 mM $NaH_2PO_4$, 0.5 mM $MgCl_2$, 25 mM Hepes, 22 mM Glucose; pH=7.4 with NaOH). Before the experiments, physiological extracellular $Ca^{2+}$ levels (1.8 mM) were gradually restored.

#### Intracellular $Ca^{2+}$ recordings

Intracellular $Ca^{2+}$ measurements were performed at 36°C. Cytosolic $Ca^{2+}$ was optically measured in ventricular CMs loaded with the probe Fluo-4 AM (10 µM) by 30 min incubation at room temperature. During the recording, cells were perfused with an extracellular solution containing (in mM): 154 NaCl, 4 KCl, 5 HEPES NaOH, 1.5 $CaCl_2$, 1 $MgCl_2$, 5.5 glucose; pH=7.35 with NaOH.

$Ca^{2+}$ transients (CaTs) were recorded as Fluo-4 signal, collected at 40X magnification (oil immersion) through a 535 nm band pass filter, converted to voltage, low-pass filtered (10 kHz) and digitized at 20 kHz. All fluorescence (F) values were normalized to $F_0$, recorded during prolonged quiescence, to compensate for potential differences in probe loading. $Ca^{2+}$ dynamics were evaluated in field-stimulated or voltage-clamped CMs, as indicated in the text.

ISTA, LEVO, and their combination were evaluated in separate groups of cardiomyocytes, with each cell serving as its own drug-free control. For combined treatment, ISTA and LEVO were applied simultaneously after baseline recordings. This design was used for both steady-state $Ca^{2+}$ measurements and the voltage-clamped protocols to limit recording duration and preserve experimental stability.

**Electrophysiological recordings**

Whole-cell patch-clamp experiments were performed in voltage-clamp mode. Cells were perfused (at 36 °C) with the extracellular solution described in the previous section. Patch pipettes (tip resistance of 1.5-2.5 MΩ) were filled with an intracellular solution containing (in mM): 110 K-aspartate, 23 KCl, 3 MgCl2, 0.04 CaCl2, 0.1 EGTA KOH ($10^{-7}$ $Ca^{2+}$-free), 5 Hepes KOH, 0.4 Na+-GTP, 5 Na+-ATP, 5 Na+-phosphocreatine; pH=7.3 with KOH. Membrane capacitance and series resistance (<10 MΩ) were measured in every cell but left uncompensated. The recorded signals were amplified by MultiClamp 700B (Molecular Devices, Sunnyvale, CA, United States), digitized at 20 kHz (Axon Digidata 1440A, Molecular Devices), and filtered at 10 kHz.

As validated in previous works [2,3], membrane current elicited by voltage steps from -40 to 0 mV in the presence of 2 mM 4-aminopyridine and 0.1 mM $BaCl_2$ are largely representative of $I_{CaL}$; intracellular $K^+$ was not substituted to avoid interference with SR function.

The change in membrane current induced by caffeine pulse (at Vm = -40 mV) was taken as a measure of $Na^+/Ca^{2+}$ exchanger current ($I_{NCX}$) [4]. To avoid $Ca^{2+}$ influx during the SR emptying, caffeine was dissolved in $Ca^{2+}$-free solution (containing 1 mM EGTA CsOH).

**Intact trabeculae**

Mice were heparinised (5000 U/kg, i.p.) and euthanized by cervical dislocation. The heart was rapidly excised and the proximal aorta was retrogradely perfused with a modified Krebs-Henseleit solution (KH) containing (mM): 120 NaCl, 15 KCl, 1.2 $NaH_2PO_4$, 2 $Mg_2SO_4$, 20 $NaHCO_3$, 1.5 $CaCl_2$, 20 butanedione-monoxime (BDM), 10 glucose, pH 7.4, equilibrated with 95% $O_2$/5% $CO_2$. High potassium concentration and BDM were important to stop spontaneous beating and to minimize contractures due to cutting damage, respectively. Thin, unbranched RV trabeculae were carefully isolated by transecting the atrioventricular ring at one extremity and the ventricular wall at the other. Trabecular diameter was measured and cross-sectional area was calculated assuming an ellipsoidal geometry. Subsequently, intact preparations were mounted between the basket-shaped end of a force transducer and a motor, in a glass chamber equipped with platinum electrodes for electrical field stimulation. The dissection solution was then replaced with a similar buffer containing low KCl (5 mM) and no BDM. The effects of drugs (ISTA, LEVO or ISTA+LEVO) were evaluated by adding them to the recirculating buffer and allowing at least 25-30 minutes for equilibration. In combination experiments, trabeculae were first exposed to ISTA; LEVO was then added while ISTA remained in the recirculating solution. Thus, the combination was compared with the preceding ISTA condition in the same preparation.

**Skinned muscles**

After heart excision, thin, unbranched, and uniform trabeculae were carefully dissected from the left ventricle. Trabeculae were incubated overnight in a cold (4°C) standard relaxing solution (containing 10 mM 3-(N-morpholino) propanesulfonic acid, 10 mM EGTA, 1 mM free $Mg^{2+}$, 5 mM MgATP, 10 mM creatine phosphate, pH adjusted to 7.0 with KOH added) with 0.5% Triton X-100, to remove cellular membrane and all cytoplasmic organelles membrane such as mitochondria and sarcoplasmic reticula. A selected skinned trabecula was attached to aluminium T-clips, so as to facilitate attachment to the setup and give preparations more stability during several measurements and mounted in the experimental setup between a force transducer and a fixed hook. Three types of solutions were employed: relaxing solution (pCa 9) with 10 mM EGTA, pre-activating solution with 0.2 mM EGTA and 9.8 mM 1,6-diamino hexane-N,N,N',N'-tetraacetic acid (HDTA), and a maximal activating (pCa 4.5) solution with 10 mM Ca-EGTA. Relaxing and maximal activating solutions were mixed in different proportions to obtain activating solutions with various pCa's. All solutions contained: 100 mM BES (N,N-bis[2-hydroxyethyl]-2-aminoethane sulphonic acid); 6.29 mM $Na_2ATP$, 6.63 mM $MgCl_2$. pH was adjusted to 7 with KOH at 20°C.

Sarcomere energetics was assessed in skinned trabeculae (21°C) by simultaneous measurement of isometric force and ATPase activity under different $Ca^{2+}$ concentrations, both in the absence and in the presence of the drugs. The force–pCa relationship was determined by exposing the preparation to solutions with pCa values ranging from 3.5 to 9.0 in a continuously stirred chamber. ATP consumption of the skinned trabeculae was measured by an enzyme-coupled assay in which NADH oxidation was monitored by measuring absorbance at 340 nm. The signal was calibrated by adding 0.5 nmol ADP, allowing ATPase activity to be calculated from the slope of the absorbance curve [5,6].

$Ca^{2+}$-activated ATPase activity was calculated by subtracting basal activity measured under relaxing conditions and was normalized to trabecular volume. Details of the experimental apparatus and the ATPase measurement principle have been described previously [7,8]. Additional experiments were conducted by exposing skinned trabeculae to increasing concentrations of LEVO in a solution containing an intermediate calcium concentration (pCa 5.8) to characterize the dose–response relationship.

### SUPPLEMENTARY FIGURES

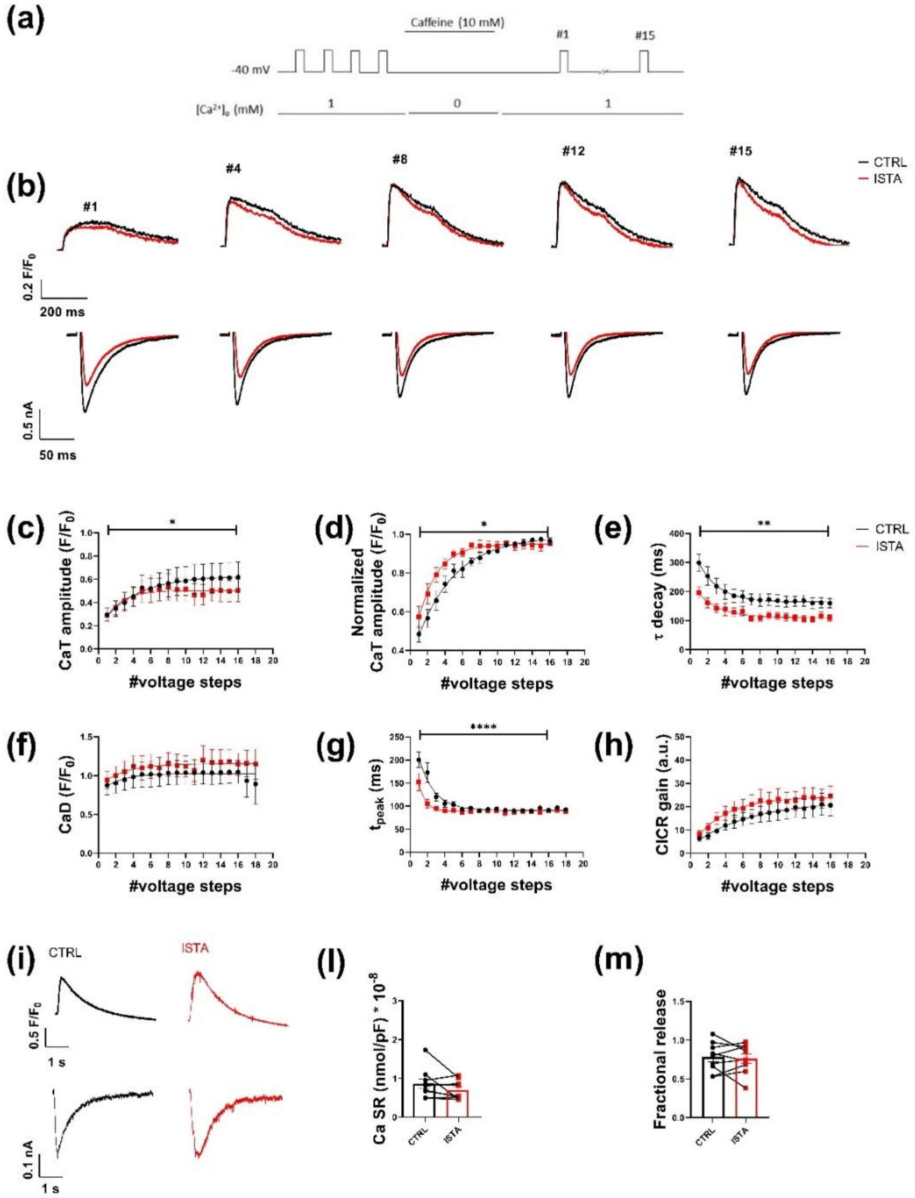


**Figure S1.** ISTA (20 µM) effect on SR $Ca^{2+}$ content and uptake. (a) Experimental protocol. (b) Representative traces of CaT (upper panels, superimposed baselines) and $I_{CaL}$ (lower panels). (c-h) Mean values of CaT parameters measured during each voltage step of the loading train. (n=9, N=2). (i) Representative traces of caffeine-evoked CaT (upper panels) and $I_m$ (lower panels). (l) Ca SR (CTRL: 0.87 ± 0.13, n=9, N=2; ISTA: 0.69 ± 0.08, n=9, N=2). (m) Fractional release (CTRL: 0.78 ± 0.06, n=9, N=2; ISTA: 0.77 ± 0.06, n=9, N=2). Data are expressed as mean ± SEM. Mixed-effect model followed by Sidak post-hoc test (panels c-h) and Wilcoxon test (panels l, m), * p<0.05, ** p<0.01, **** p<0.0001.

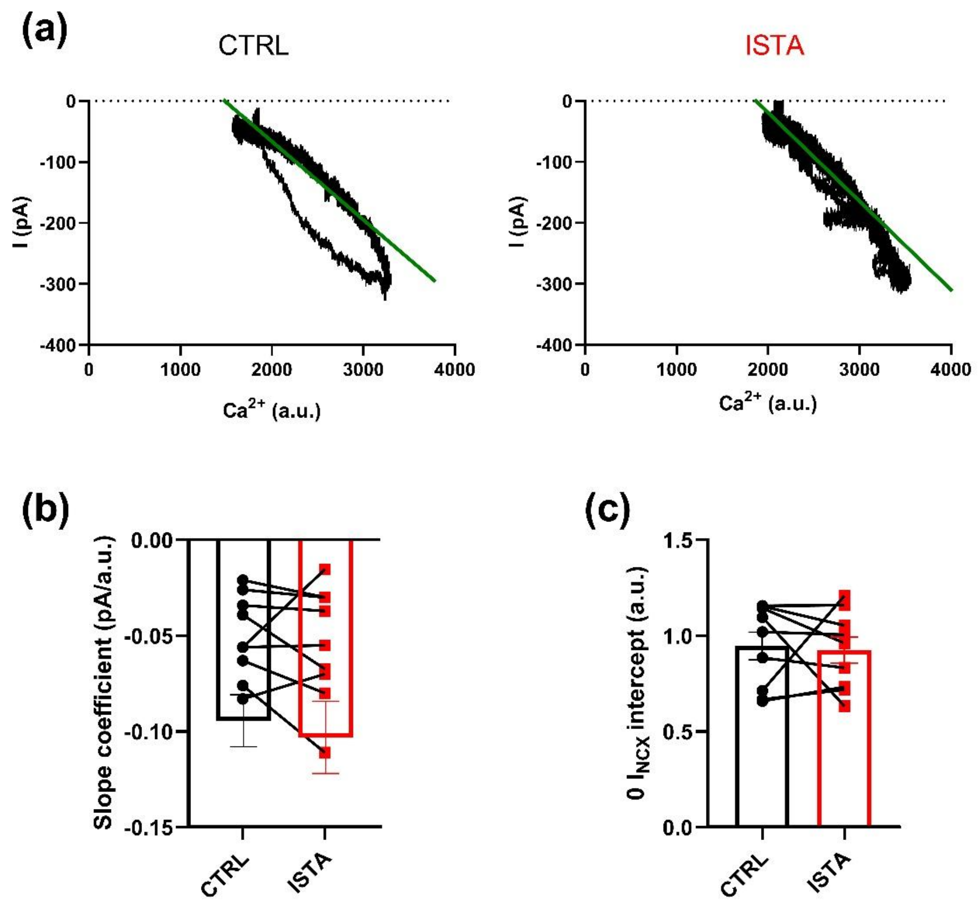


**Figure S2.** ISTA effect on the relationship between $I_{NCX}$ and [$Ca^{2+}$]. (a) Representative $I_{NCX}$/[$Ca^{2+}$] loops with linear regression (green lines). (b) Line slope coefficient (CTRL: -0.05 ± 0.01, n=9, N=2; ISTA: -0.06 ±0.01, n=9, N=2). (c) 0 $I_{NCX}$ intercept (CTRL: 0.94 ± 0.07, n=9, N=2; ISTA: 0.92 ± 0.07, n=9, N=2). Data are expressed as mean ± SEM. Paired Student's t-test.

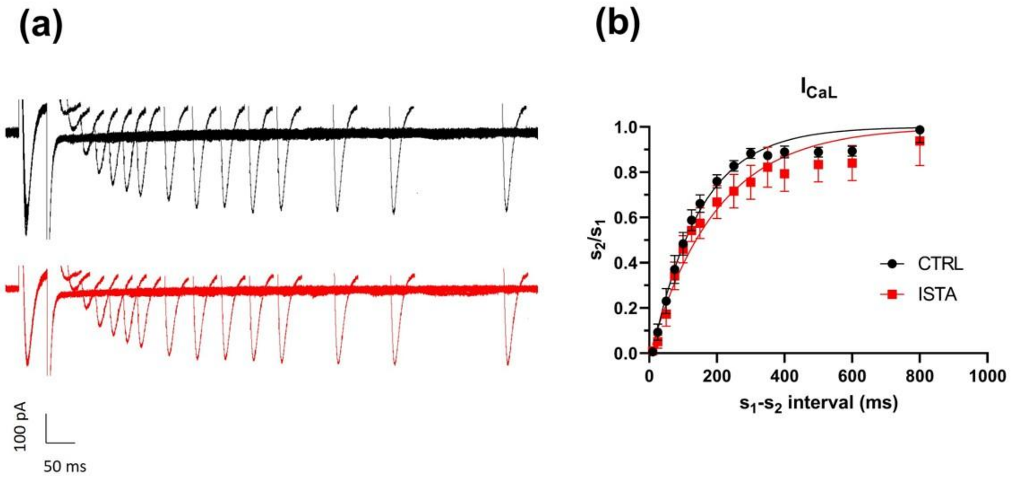


**Figure S3.** ISTA effect on $I_{CaL}$ restitution. (a) Representative traces of $I_{CaL}$ recorded during the restitution protocol. (b) Restitution curves. (n=14, N=2). Data are expressed as mean ± SEM. Mixed-effect model followed by Sidak post-hoc test.

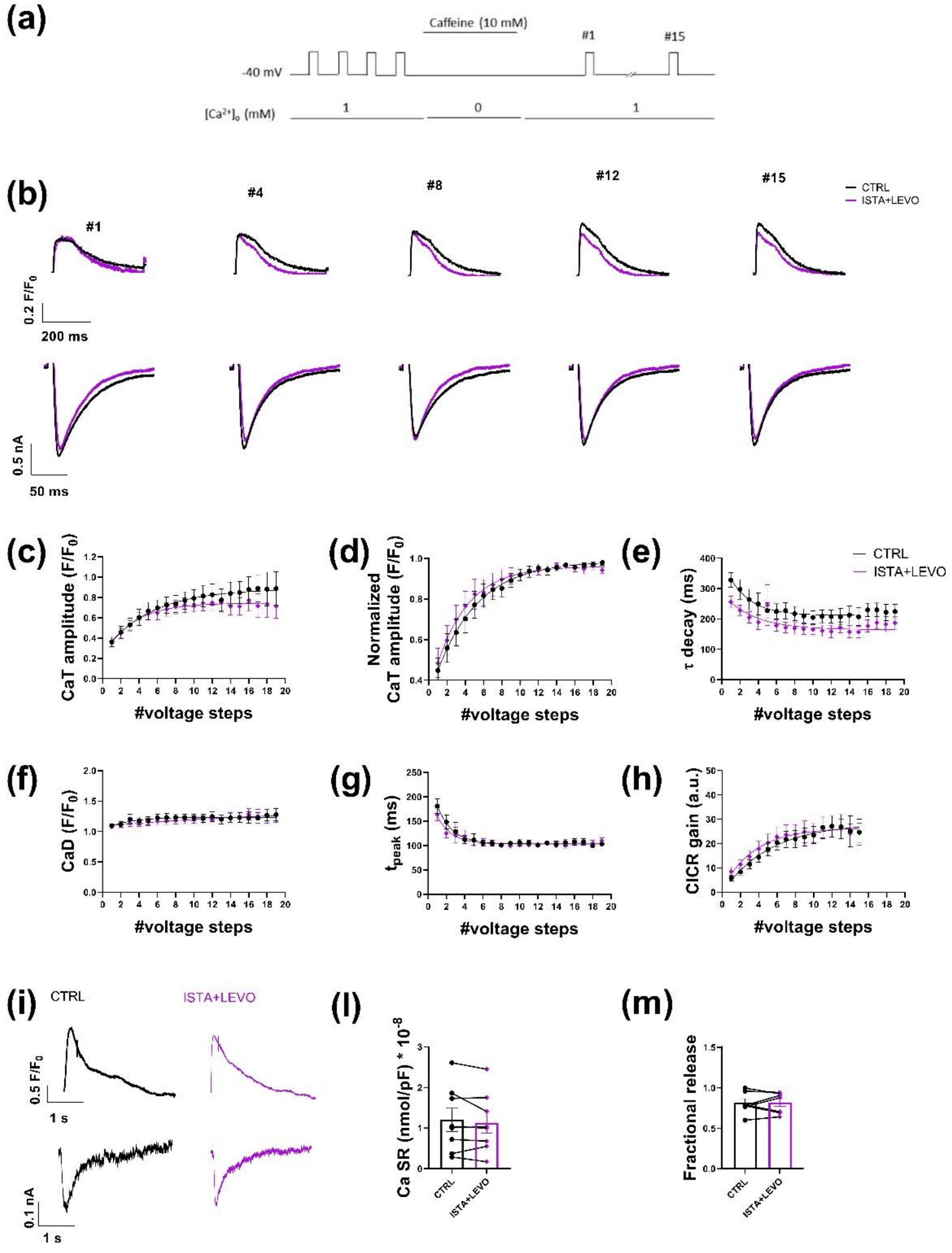


**Figure S4.** ISTA+LEVO effect on SR $Ca^{2+}$ content and uptake. (a) Experimental protocol. (b) Representative traces of CaT (upper panels, superimposed baselines) and $I_{CaL}$ (lower panels). (c-h) Mean values of CaT parameters. (n=10, N=2). (i) Representative traces of caffeine-evoked CaT (upper panels) and $I_m$ (lower panels). (l) Ca SR (CTRL: 1.20 ± 0.28, n=8, N=2; ISTA+LEVO: 1.13 ± 0.26, n=8, N=2). (m) Fractional release (CTRL: 0.81 ± 0.05, n=7, N=2; ISTA+LEVO: 0.081 ± 0.05, n=7, N=2). Data are expressed as mean ± SEM. Mixed-effect model followed by Sidak post-hoc test (panels c-h) and Student's t-test (panels l, m).

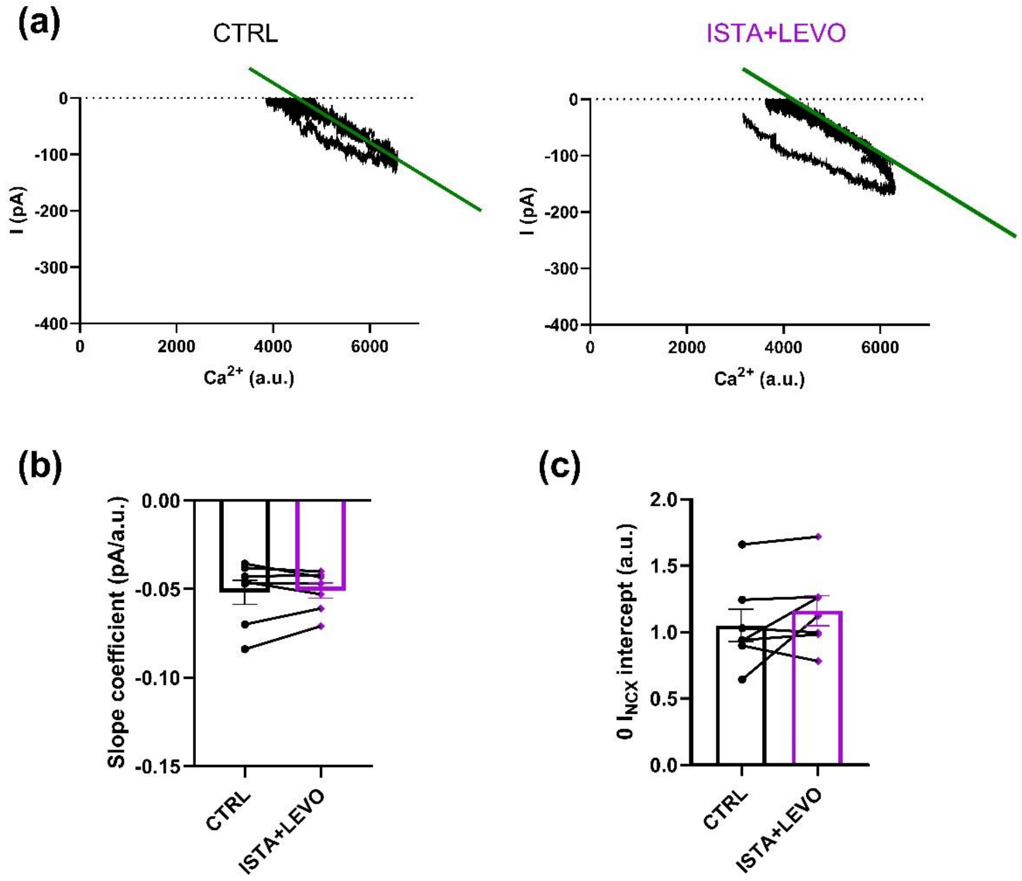


**Figure S5.** ISTA+LEVO effect on the relationship between $I_{NCX}$ and $[Ca^{2+}]$. (a) Representative $I_{NCX}/[Ca^{2+}]$ loops with linear regression (green lines). (b) Line slope coefficient (CTRL: -0.05 ± 0.007, n=7, N=2; ISTA+LEVO: -0.05 ± 0.004, n=7, N=2). (c) 0 $I_{NCX}$ intercept (CTRL: 1.049 ± 0.122, n=7, N=2; ISTA+LEVO: 1.161 ± 0.113, n=7, N=2). Data are expressed as mean ± SEM. Paired Student's t-test.

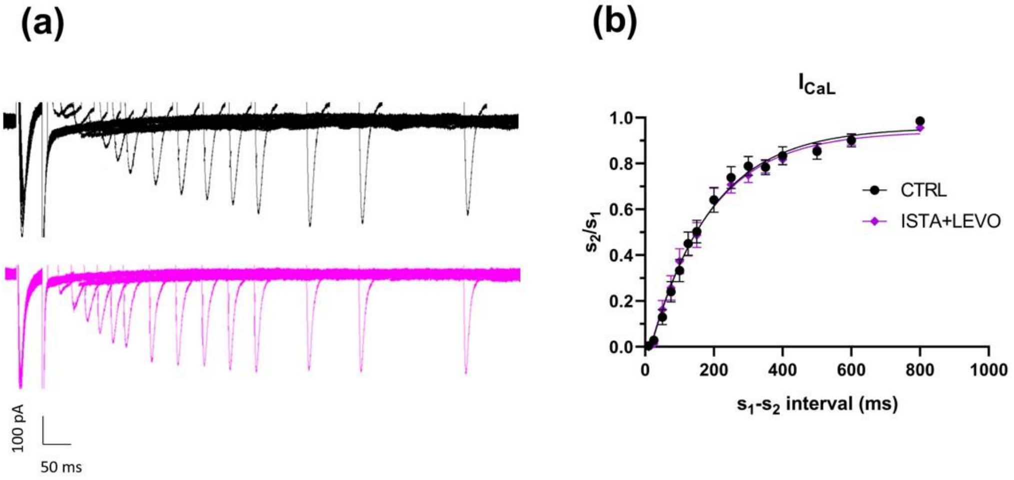


**Figure S6.** ISTA+LEVO effect on $I_{CaL}$ restitution. (a) Representative traces of $I_{CaL}$ recorded during the restitution protocol. (b) Restitution curves. (n=9, N=1). Data are expressed as mean ± SEM. Mixed-effect model followed by Sidak post-hoc test.

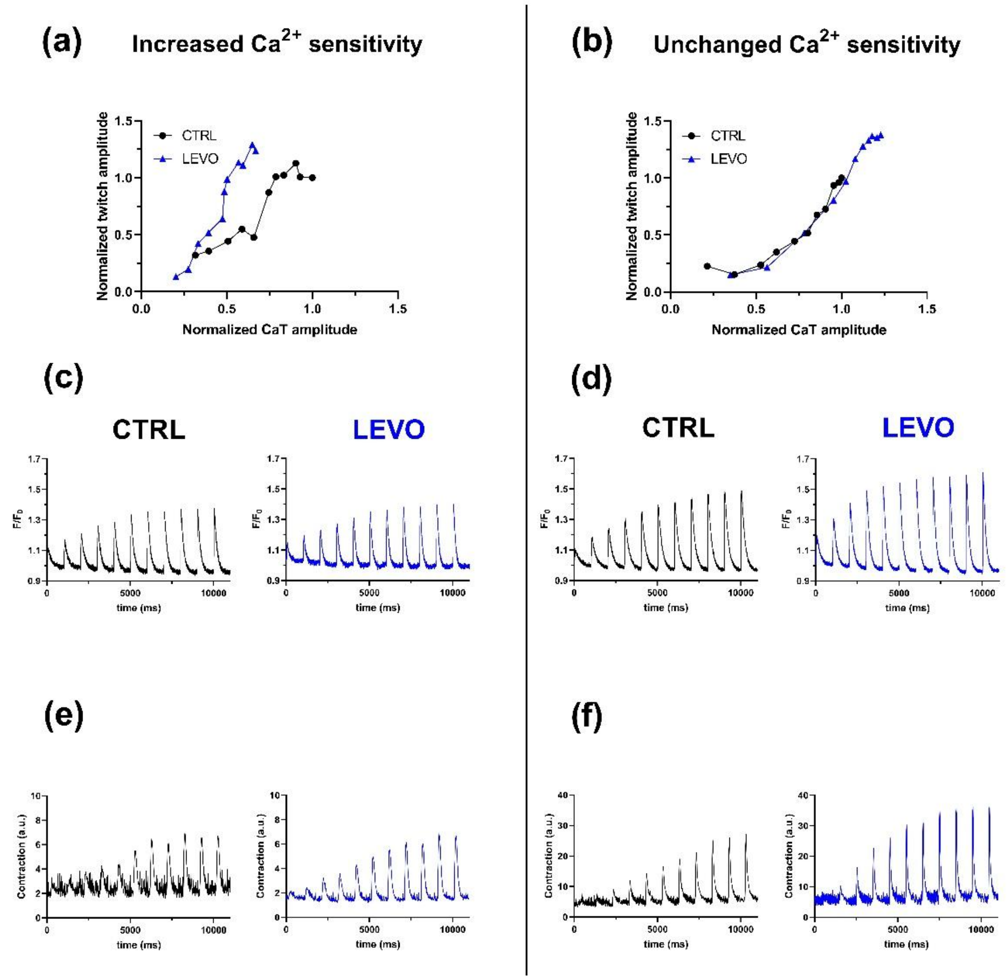


**Figure S7.** Effects of LEVO on contractile performance in isolated cardiomyocytes. (a, b) Representative twitch contraction/CaT plots (normalized to the corresponding steady-state CTRL values) from a responding cell (showing increased $Ca^{2+}$ sensitivity after LEVO perfusion) and a non-responding cell (showing unchanged $Ca^{2+}$ sensitivity), respectively. (c, d) Representative CaT traces elicited by depolarizing voltage steps of increasing amplitude, in CTRL and after LEVO perfusion. (e, f) Representative twitch contraction traces recorded with the same protocol.

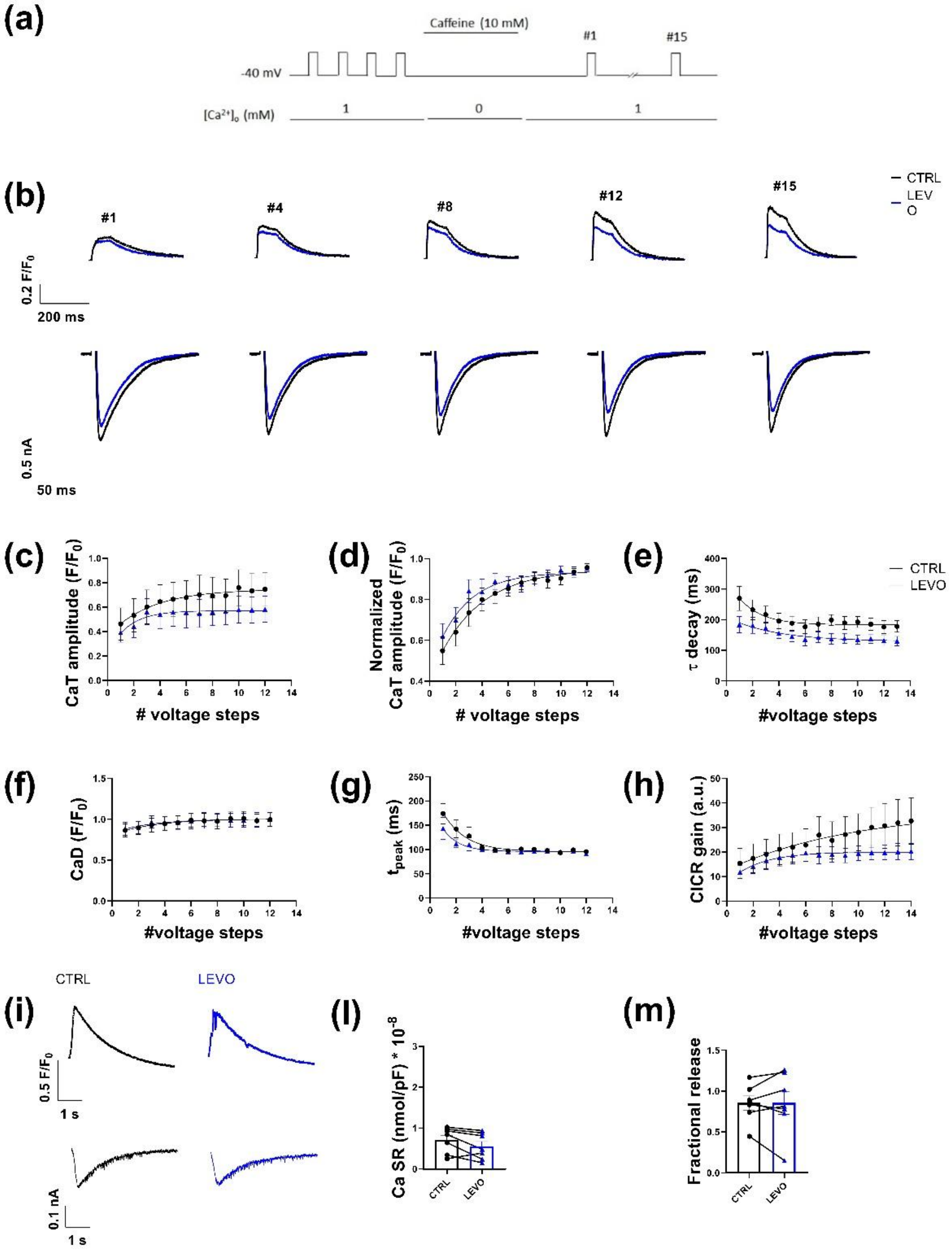


**Figure S8.** LEVO effect on SR $Ca^{2+}$ content and uptake. (a) Experimental protocol. (b) Representative traces of CaT (upper panels, superimposed baselines) and $I_{CaL}$ (lower panels). (c-h) Mean values of CaT parameters. (n=9, N=2). (i) Representative traces of caffeine-evoked CaT (upper panels) and $I_m$ (lower panels). (l) Ca SR (CTRL: 0.74 ± 0.11, n=8, N=2; LEVO: 0.51 ± 0.11, n=8, N=2). (m) Fractional release (CTRL: 0.85 ± 0.09, n=7, N=2; LEVO: 0.85 ± 0.14, n=7, N=2). Data are expressed as mean ± SEM. Mixed-effect model followed by Sidak post-hoc test (panels c-h) and Student's t-test (panels l, m).

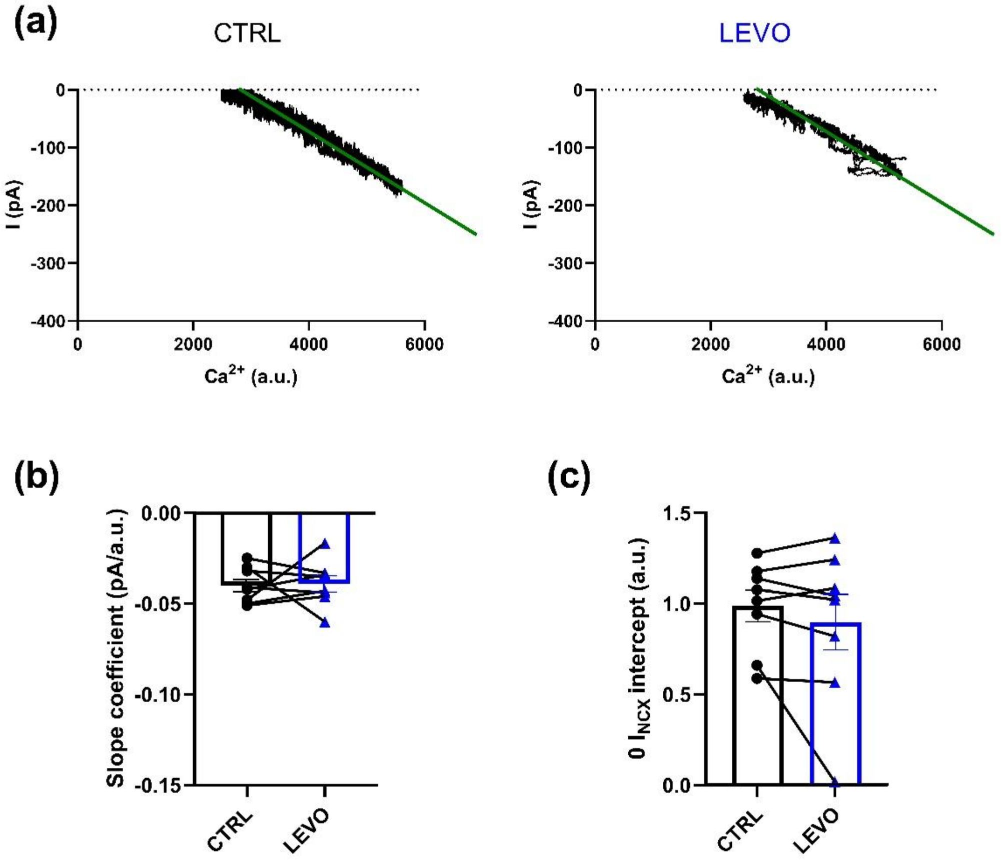


**Figure S9.** LEVO effect on the relationship between $I_{NCX}$ and $[Ca^{2+}]$. (a) Representative $I_{NCX}/[Ca^{2+}]$ loop with linear regression. (b) Line slope coefficient (CTRL: -0.04 ± 0.003, n=8, N=2; LEVO: -0.04 ± 0.004, n=8, N=2). (c) 0 $I_{NCX}$ intercept (CTRL: 0.99 ± 0.09, n=8, N=2; LEVO: 0.90 ± 0.015, n=8, N=2). Data are expressed as mean ± SEM.

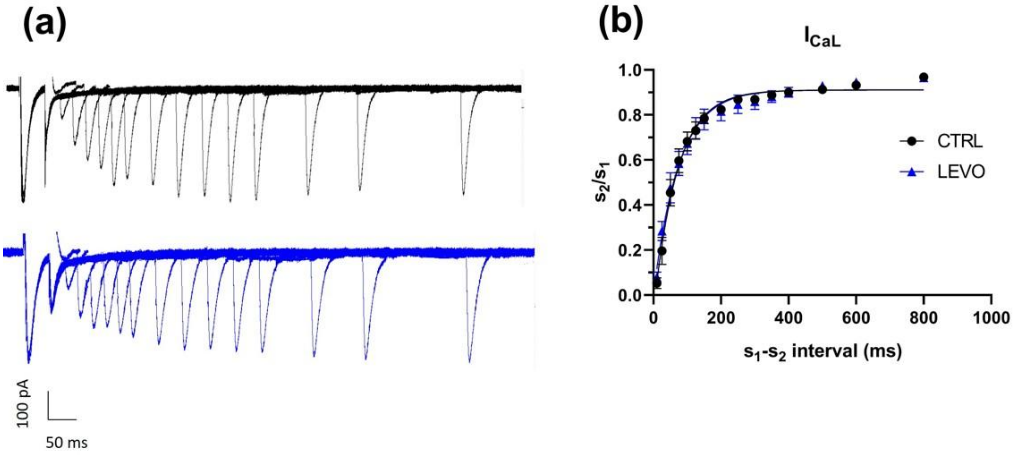


**Figure S10.** LEVO effect on $I_{CaL}$ restitution. (a) Representative traces of $I_{CaL}$ recorded during the restitution protocol. (b) Restitution curves. (n=10, N=3). Data are expressed as mean ± SEM, Mixed-effect model followed by Sidak post-hoc test.
